# bcGST - an interactive bias-correction method to identify over-represented gene-sets in boutique arrays

**DOI:** 10.1101/240234

**Authors:** Kevin YX Wang, Alexander M Menzies, Ines P Silva, James S Wilmott, Yibing Yan, Matthew Wongchenko, Richard F Kefford, Richard A Scolyer, Georgina V Long, Garth Tarr, Samuel Mueller, Jean YH Yang

## Abstract

**Motivation:** Gene annotation and pathway databases such as Gene Ontology and Kyoto Encyclopedia of Genes and Genomes are important tools in Gene Set Test (GST) that describe gene biological functions and associated pathways. GST aims to establish an association relationship between a gene set of interest and an annotation. Importantly, GST tests for over-representation of genes in an annotation term. One implicit assumption of GST is that the gene expression platform captures the complete or a very large proportion of the genome. However, this assumption is neither satisfied for the increasingly popular boutique array nor the custom designed gene expression profiling platform. Specifically, conventional GST is no longer appropriate due to the gene set selection bias induced during the construction of these platforms.

**Results:** We propose bcGST, a bias-corrected Gene Set Test by introducing bias correction terms in the contingency table needed for calculating the Fisher’s Exact Test (FET). The adjustment method works by estimating the proportion of genes captured on the array with respect to the genome in order to assist filtration of annotation terms that would otherwise be falsely included or excluded. We illustrate the practicality of bcGST and its stability through multiple differential gene expression analyses in melanoma and TCGA cancer studies.

**Availability:** The bcGST method is made available as a Shiny web application at http://shiny.maths.usyd.edu.au/bcGST/

## 1 Introduction

Gene expression profiling platforms have enabled researchers to explore complex human diseases on an ever larger scale. As our knowledge of disease mechanisms increases, there is a greater need for more targeted and reproducible analysis requiring ever more sensitive expression platforms for clinical biomarkers. A boutique array platform is well-suited to tackle these challenges because the built-in probes are specially designed for the disease of interest. Depending on manufacturer designs, these probes can measure signals with greater sensitivity than other platforms such as microarray and RNA-Seq. This increase in sensitivity is typically achieved by reducing the number of probes to those directly relevant to the disease of interest, which increases the noise-to-signal ratio during competitive hybridisation.

However, this reduction in the number of probes also brings new challenges to traditional pre-processing tools and statistical analyses (Jung and Sohn, 2014). Specifically, many bioinformatics and statistical techniques have been developed for the purpose of biological discovery and these are not appropriate for targeted platforms like boutique arrays. One such example is the gene set over-representation test, commonly referred to as Gene Set Test (GST), originally developed for microarray experiments (Subramanian *et al*., 2005; Irizarry *et al*., 2009). In GST, instead of interpreting genes directly, these genes are mapped to biologically meaningful annotations in curated biological databases. Then, by applying appropriate statistical tests, statistical significance is assigned to the annotation terms instead of specific genes. Typically, genes that are differentially expressed (DE) are of particular interest in GST because these genes are potential biomarkers which could inform us of the underlying disease mechanisms if their association with biological functions and pathways can be established.

When performing GST on boutique arrays, one often ignored issue is that the probes built onto the gene expression profiling platform do not necessarily match with the gene set database that is of interest. For example, most GST databases, such as Gene Ontology (GO) (Consortium, 2000), were built with the whole human genome in mind. As a result, the intended gene universe when performing GST is the whole genome, or at least, a genome-wide gene expression profiling platform. The particular challenge for boutique arrays is that boutique array genes are specialised by design, and thus in much smaller quantity (ranging from 100 to a maximum of 1,000 probes) compared to the whole genome. Statistical testing of annotation terms is therefore restricted to genes on the boutique arrays. We refer to this issue as gene set selection bias. While there was a tremendous amount of research into making evermore advanced methods and accessible web-based tools (Subramanian *et al*., 2005; Backes *et al*., 2007; Zheng and Wang, 2008; Eden *et al*., 2009; Wu *et al*., 2010), there is always an implicit assumption that end-users use a large gene expression profiling platform in experiments. In light of this, there is a need to develop a GST methodology which can produce interpretable gene set enrichment analysis results in the presence of gene set selection bias.

This paper presents a bias-corrected GST, abbreviated as bcGST, a novel method that enables the application of GST in the absence of large parts of the genome, as we typically encounter in boutique arrays. Using a published microarray and TCGA RNA-Seq data, we create synthetic simulations of boutique array simulation to examine the advantages and practicality of bcGST. The bcGST method corrects for the gene set selection bias effect by reducing false negatives during scoring gene set significance. We further demonstrate bcGST's ability to yield stable gene set test results in the presence of gene set selection bias. Application of bcGST to a real boutique array study and a TCGA RNA-Seq data revealed key cancer pathways despite low concordance between the two platforms. We also developed a Shiny (Chang *et al*., 2017) web application for the bcGST method. This application enables the detection and exploration of genesets through interactive visualisation and adjustments.

## 2 Materials and Methods

### 2.1 Datasets

#### 2.1.1 Microarray melanoma data

A published gene expression study (Schramm *et al*., 2013) from stage III melanoma patients was used to illustrate our methods. The platform used was Illumina Human WG-6 BeadChip microarray, version 3. In this study, a good prognosis group (*n =* 25) was defined as those samples with more than four years survival with no sign of relapse and a poor prognosis group (*n* = 22) as those samples that died within one year of metastasis. Raw data were processed using the NEQC method (Shi *et al*., 2010). Negative control probes were used for background correction and quantile normalisation used both negative and positive probes. The normalised data has 23,460 probes, corresponding to 17,934 unique gene symbols.

#### 2.1.2 TCGA cancer gene expression data

The Cancer Genome Atlas (TCGA) RNA-Seq expression and clinical information (Weinstein *et al*., 2013) were downloaded using the R (R Development Core Team, 2017) package `ExperimentHub` (Maintainer, 2016) from Gene Expression Omnibus (GEO) with submission ID GSE62944, on 11 October 2016. The voom method available in the `limma` (Ritchie *et al*., 2015) package was used to normalise RNA-Seq count data so the processed data could be analysed using available microarray analysis methodologies (Law *et al*., 2014).

There were 23,368 genes across all 19 cancers. We used this pan-cancer data in two ways. First, for the purpose of creating simulated boutique array experiments, patients with either tumour status or vital status missing were discarded. Differential gene expression analysis was performed between good prognosis patients who were both tumour-free and alive, against poor prognosis patients who carried tumours and died. Under this classification, cancers with less than 10 samples in either prognosis groups were eliminated. Second, for the purpose of demonstrating our method on a real boutique array dataset, we selected from the TCGA data the Skin Cutaneous Melanoma (SKCM) patients with American Joint Committee on Cancer (AJCC) tumour stage classification.

#### 2.1.3 NanoString customised panel

This is a customised gene expression profiling panel containing 800 genes designed to cover a broad spectrum of melanoma biology, immune-related genes, cellular functions, and signalling pathway transcriptional targets. See Table S3 for a complete list of genes. Similar to other boutique array panels, the manufacturer argues that due to gene co-expression, a well-designed boutique array can measure gene expression with a smaller set of probes and is expected to capture a significant amount of expression variability (NanoString Technologies, 2015).

Due to the non-negligible differences between how microarrays, RNA-Seq and NanoString collect signals from their respective panels (Robinson *et al*., 2015; Chen *et al*., 2016; Nookaew *et al*., 2012; Guo *et al*., 2013), we used this panel in two different ways. First, for the purpose of creating simulated boutique array experiments, we are only interested in the induced biological knowledge that the NanoString genes bring to the analysis. Hence in all simulations, we only used the NanoString gene symbols to subset genes from the larger platforms, namely the microarray and RNA-Seq data described above. Second, we used the gene expression profiles of 95 samples from a study of the effect of targeted therapy and immunotherapy for BRAF-mutant patients in melanoma (Silva *et al*., 2017). Normalisation was performed using the `NanoStringQCPro` package (Nickles D *et al*., 2017) with content normalisation.

### 2.2 Simulated boutique array experiments

Given the microarray and TCGA RNA-Seq data, we took the subset of NanoString PanCancer panel genes to construct smaller datasets which resemble data collected from boutique array experiments. We will refer to these sub-datasets as “simulated boutique arrays” or just “simulations" because they contain gene expression measurements for specialised gene sets without actually performing the experiments. Such simulations allowed for a higher degree of control for our proposed correction method and eliminate possible experimental inconsistencies. No further normalisation or data processing was performed on the simulations. During the evaluation, the original inference results from the larger platforms (i.e. microarray and TCGA RNA-Seq) were treated as the gold standard against which inference results from the simulations are compared.

The first experiment was simulated with respect to the Stage III microarray, in which there were 663 genes common to both platforms. The second experiment was simulated with respect to the TCGA RNA-Seq data, with 775 genes common to both platforms, across all cancer types. All samples were retained during construction of the simulations.

#### 2.2.1 Differential expression and GO analysis

We used empirical Bayes moderated t-test statistics (Smyth, 2004) implemented in the `limma` package from R (R Development Core Team, 2017) to perform differential gene expression analysis between patient groups in each dataset.

In the simulation studies, microarray experiment genes with a p-value less than 0.01 were considered as differentially expressed (DE). For the TCGA data, genes with a p-value less than 0.005 were considered as DE. These thresholds were chosen to yield a reasonable number of DE genes so that we have a reasonable mapping of annotation terms. We further discarded TCGA cancer datasets with fewer than 5 differentially expressed genes on the simulated boutique array, the full list of which are on Table S1a.

In the real data application, differential gene expression analysis was performed between patients who were classified as AJCC Stage IVC and those samples in either Stage IVA, IVB or IIIC (pooled). We used a less conservative p-value of 0.05 on both the NanoString and the TCGA SKCM RNA-Seq data as a cut-off for a gene to be considered DE. Table S1b shows a list of samples.

In the simulation study, DE genes were then mapped to the GO database via Entrez ID. Enrichment tests and further parameter extractions were all performed through `topGO` package (Alexa, 2015). We considered only the ontology branch “Biological Processes” of the GO database, keeping only terms with at least 10 annotated genes, and further discarded all GO terms with missing or undefined parameters from our analysis. In the real data application, we used the BioCarta pathway downloaded from the Molecular Signatures Database from Broad Institute, version 6.0 (Subramanian *et al*., 2005; Liberzon *et al*., 2011).

### 2.3 Gene Set Enrichment Analysis

#### 2.3.1 Fisher’s Exact Test parameters

We used Fisher’s Exact Test (FET) to test for over-representation of differentially expressed (DE) genes with the GO annotation terms. Under the null hypothesis that relative proportions of one gene set are independent of a second gene set, parameters on a two-by-two contingency table follow a hypergeometric distribution, so a p-value can be calculated based on this distribution.

To clarify the relationships between boutique array genes, differentially expressed genes and genes assigned to pathways, we have tabulated the number of genes in each category in Table 1a and Table 1b.

**Table 1a.** Contingency table of boutique array genes, split according to if these are differentially expressed and assigned to a given pathway.

| Boutique array gene universe | DE | not DE |
| --- | --- | --- |
| Pathway | $a$ | $b$ |
| Not pathway | $e$ | $f$ |

**Table 1b.** Contingency table of non-boutique array genes, split according to if these are differentially expressed and assigned to a given pathway.

| Non-boutique array gene universe | DE | not DE |
| --- | --- | --- |
| Pathway | $c$ | $d$ |
| Not pathway | $g$ | $h$ |

In a FET, we refer to all genes under consideration as the “gene universe”. Depending on the gene expression profiling platform being used, the gene universe will change accordingly. For example, if we wish to make inference only within a boutique array, then we simply take the gene universe to be the boutique array. Equation (1) computes the onesided FET p-value, where numbers (*a, b*, *e* and *f*) are defined in Table 1a, for each pathway under consideration. If the whole genome is the gene universe, then the p-value is calculated using Equation (2), where the numbers *(a* + *c*, *b* + *d, e* + *g* and *f* + *h*) are shown in Table 1c. As most curated biological annotation databases like GO and Kyoto Encyclopedia of Genes and Genomes (KEGG) were developed with respect to the whole genome, it is therefore important to know which gene universe is under consideration. Misspecifying the gene universe as the boutique array genes when the whole genome is intended can induce gene set selection bias into GST, see Figure 1.

**Table 1c.** Contingency table of all genes in a genome-wide gene universe, split according to if these are differentially expressed and assigned to a given pathway.

| Genome-wide gene universe | DE | not DE |
| --- | --- | --- |
| Pathway | $a + c$ | $b + d$ |
| Not pathway | $e + g$ | $f + h$ |

**Fig. 1.**
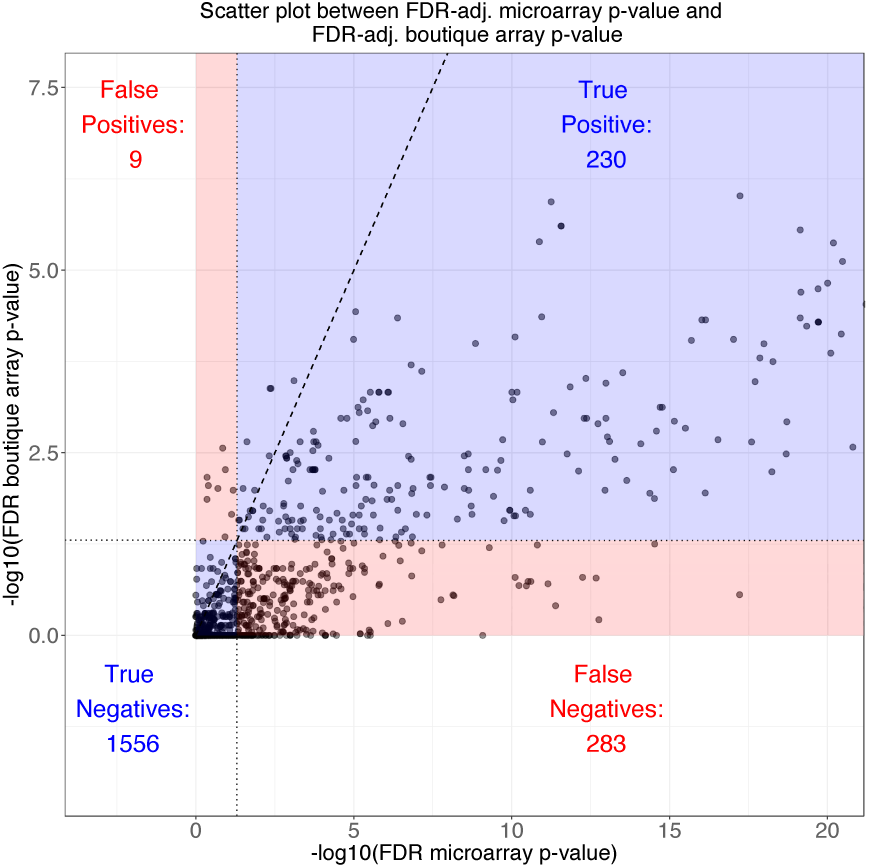
A scatter plot of 2,078 Gene Ontology (GO) terms common to both the microarray data and the simulated boutique array data. The *x*- and *y*-axes are the FDR-adjusted FET p-values from the microarray and simulated boutique array on a − log_10_ scale, respectively. 1,519 out of 2,078 (73%) points are below the line of no change (*y* = *x*, black dashed line). By dividing the figure at FDR-adjusted p-values < 0.05 for both gene universes, we can further confirm the majority of false decisions made by FET p-values came from false negatives.

Equation (1) is the standard way of calculating over-representation p-value using FET Here, the hypothesis is the DE gene list is not associated with the pathway under consideration. The competitive alternative hypothesis is that there is an over-representation of DE genes in the pathway, hence we formulated a one-sided test. In Table 1a, *a* is the number of differentially expressed genes among all genes on the pathway in a boutique array, and *b*, *c*,…, *h* are defined similarly by reading their corresponding column and row label in Table 1a and Table 1b.

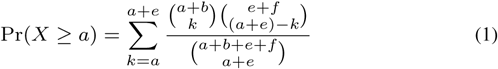

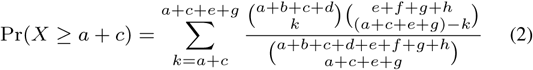

### 2.4 FET p-value grid and bcGST

In a boutique array experiment, for each pathway, parameters *a* and *e* in Table 1a and 1c are known while parameters *c* and *g* in Table 1b and 1c are unknown. We denote 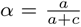 and 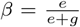 to represent different proportion of DE genes built onto the boutique array (see Results for more detail on the interpretation of these two parameters). If the value of *α* and *β* is known, then we can compute estimates (rounded to the nearest integer if necessary) of the unobserved *c* and g parameters through *ĉ* = *a*(1/*α* − 1) and *ĝ* = *e*(1/*β* − 1), respectively. The number of DE genes can then be estimated with *a* + *ĉ* + *e* + *ĝ* and the value *a* + *b* + *c* + *d* can be obtained from curated pathway database, thus completing the contingency table in Table 1c. In the process of doing so, we calculated a FET p-value by taking into account the missing genes on the boutique array, and thus this process would allow us to correct for the gene set selection bias in a boutique array GST.

However, when performing a real boutique array experiment, the precise value of *α* and *β* is never known. We thus propose a visualisation-inspired technique to evaluate the FET p-value over a grid of *α* and *β* values. The rationale behind this grid construction is that a higher value of *α* and *β* means more DE genes was built onto the boutique array, these proportions represent the severity of gene set selection bias for each pathway term. If the p-value associated with a pathway is always significant regardless of *α* and *β*, then we can conclude its initial statistical significance was not likely to be driven by these proportions and thus the gene set selection bias. Thus, we may conclude the statistical significance is robust against this bias.

To perform this computation, we will first define a range of plausible values for *α* and *β*, which can be as large as [0, 1] × [0,1], where either parameter can take any value between 0 and 1. For each ( *α*, *β*) pair on this grid, we can calculate unknown FET contingency table parameters as described above, thus obtaining a grid of FET p-values.

To simplify interpretations, we propose to further summarise the FET p-value grid into a single statistic. For each pathway or annotated term, a natural way to condense this grid information is to count the number of times which the p-value fall below a pre-determined significance threshold. This defines a grid count statistic, and we can thus consider a pathway to be robust against the induced gene set selection bias if this count statistic is high when compared to the size of the *αβ* grid. The bcGST method we are proposing is consist of both the grid of p-values and the grid count statistic.

## 3 Results

### 3.1 Gene-set selection bias presents a unique challenge to GST in boutique array experiments

Gene set selection bias is a particularly challenging problem for boutique arrays. A large proportion of genes and their associated pathways/annotation terms are left out due to the design of the boutique array and so the parameters in Table 1b will be completely ignored in the analysis. Statistical significance of annotation terms is therefore biased towards the genes on the boutique arrays due to selection bias. This bias is present even in the case of well-designed boutique arrays. See the Materials and Methods section for the detailed statistical formulation of the selection bias problem.

Figure 1 shows a scatter plot comparing FET p-values from the Stage III melanoma microarray experiment and simulated NanoString boutique array for 2,078 common GO terms. Here we regard the microarray p-values as the “truth” against which we compare the simulated boutique array p-values. The simulated boutique array FET provided correct decisions for most GO terms. There is strong monotonicity between the p-values derived from the two gene universes as we should expect from a specialised cancer gene panel. While a change in statistical significance was anticipated, 1,519 (73%) GO terms exhibited a larger p-value. Furthermore, the majority of false decisions originated from false negatives; that is, GO terms which were significant in the underlying microarray, but failed to be judged as significant in the simulation. Our bcGST method addresses this issue of gene-set selection bias.

### 3.2 P-value correction is needed to account for selection bias in boutique arrays

In practice, depending on the gene expression profiling platform being used, its gene universe should change accordingly. For example, in an Affymetrix array, the built gene-set covers the whole genome most of the time. However, if we wish to analyse with a boutique array, then the gene universe for the boutique array is much smaller in comparison. In reference to Table 1a, the gene universe has a total of *a* + *b* + *c* + *d*, the sum of all the parameters in the contingency table. In the case of the NanoString platform, the collection of genes is at most 800 genes. Performing a GST directly in the latter case will result in biased p-value calculations. To this end, we developed a correction method to account for such selection bias.

The bcGST method relies on two key ratios. For a typical gene set of interest, say an annotation term in GO, the first ratio *α* represents the proportion of DE genes on the boutique array compared to all DE genes in the genome present in the pathway. Similarly, the second ratio *β* represents the proportion of DE genes on the boutique array compared to all DE genes in the genome absent from the pathway. Using the notations from Table 1a and 1b, 

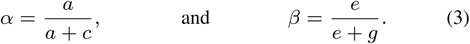

For example looking at GO:0023014, signal transduction by protein phosphorylation, had a p-value of 0.24 in the simulation above. Let *α* and *β* take values, 0.35 and 0.10 respectively; that is, the boutique array captures 35% and 10% of DE genes present and absent from the microarray. Then we can find *ĉ* = *a*(1/*α* − 1) and *ĝ = e*(1/*β* − 1), which are estimates of the unobserved parameters *c* and *g* in a typical boutique array experiment. These estimates allow us to complete a FET contingency table. Thus, a plausible bias-corrected p-value from our GST analysis is 0.003 instead. Note that this correction depends strongly on the selected (assumed but unknown) values of *α* and *β*.

### 3.3 The bcGST method corrects for false negatives for unknown *α* and *β*

We propose to use bcGST, a three-step-grid-based approach to examine the statistical significance of annotation terms under a range of *α* and *β* values. First, we consider a range of *α* and *β* values, thus constructing a grid. Second, for each pair of ratios, we complete the contingency table and perform FET for each *(α, β)* pair. This yields a grid of p-values. Third, by counting the number of p-values that fall below a certain level of statistical significance, we construct a count statistic which can better assess the statistical significance of annotation terms. We use the bcGST approach to correct for statistical significance as a result of gene-set selection bias and we will refer to the grid count statistic as the “bcGST statistic".

Figure 2 shows a heatmap of p-values for GO:0044711 (single-organism biosynthetic process) and GO:0023014 (signal transduction by protein phosphorylation). Under a conventional significance level of 0.05, both of these terms would be judged as non-significant. If we use a equally spaced *(α, β)* grid of size 400 to compute the grid count statistic, that is, counting the number of grid points that have an associated FET p-value < 0.05, we obtain bcGST statistic value of 157 for GO:0044711 and 353 for GO:0023014, respectively. Since a comprehensive range of *α* and *β* values were considered, the almost two-fold difference in the bcGST statistics between these two GO terms provides a good indication that the latter is more likely to be considered significant in a genome-wide microarray. This is indeed the case, the microarray p-value for G0:0044711 is 0.230 and the p-value for G0:0023014 is 0.005. This is one example where our bcGST statistic successfully prevents false negative conclusion (for G0:0023014).

**Fig. 2.**
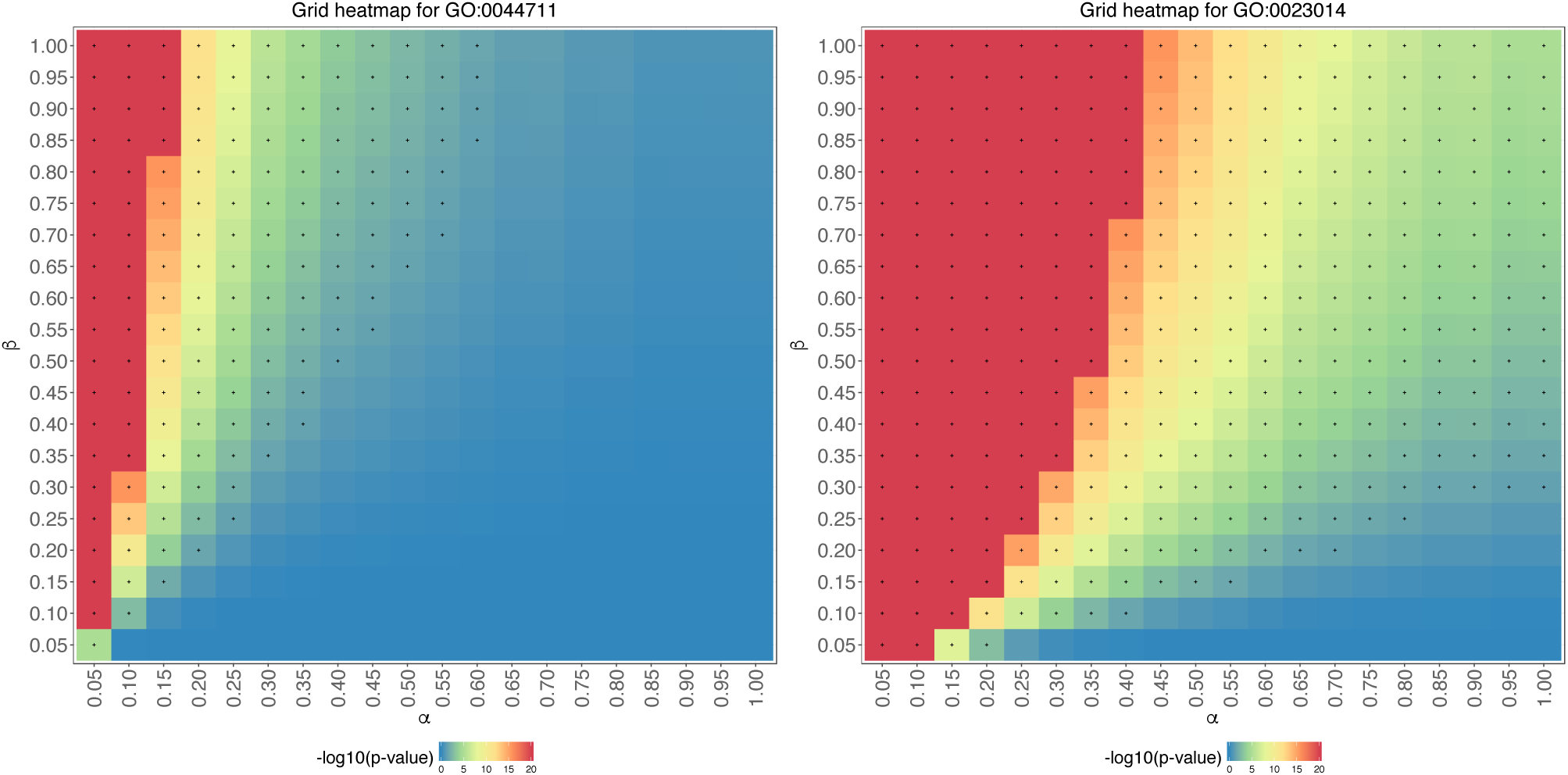
A pair of GO terms with similar simulated boutique array FET p-values, but noticeably different microarray FET p-values and bcGST statistics. The heatmaps are on a − logio scale, capped at 20 for ease of reading. Grid points with a p-value less than 0.05 are marked with a “+" symbol. (Left) GO:0044711, single-organism biosynthetic process, microarray p-value = 0.23, simulated boutique array p-value = 0.43, bcGST statistics = 157/400. (Right) GO:0023014, signal transduction by protein phosphorylation, microarray p-value = 0.005, simulated boutique array p-value = 0.24, bcGST statistics = 353/400.

Figure 3 is similar to Figure 1 but all GO terms are now coloured by grouping the bcGST statistic into four low-count/high-count categories. We note that 82% of false negatives received moderately high (> 300 grid points) or very high counts (> 350 grid points). The high bcGST statistics provide evidence for these GO terms from being incorrectly judged as nonsignificant. Thus, this count statistic supplements the boutique array FET results in overcoming the problem of false negatives caused by gene-set selection bias.

**Fig. 3.**
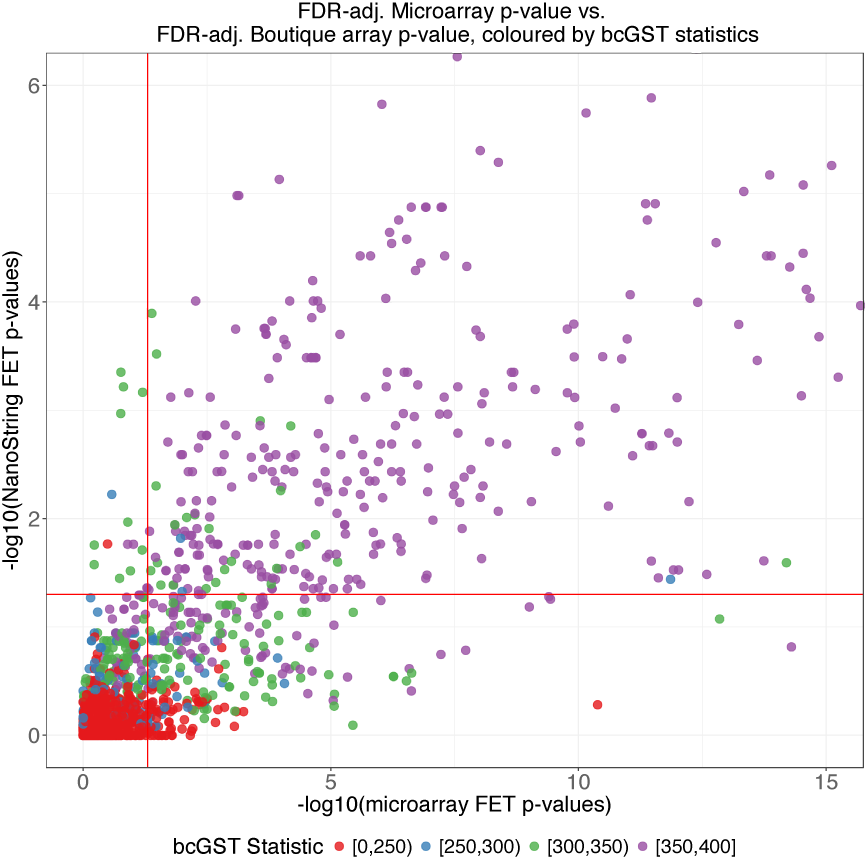
Colouring the previous scatter plot with four categories of low to high grid count statistics. We have significantly increased detections of GO terms in the false negative (lower right) region; 244 out of 283 (86%) GO terms had bcGST statistics higher than 300 out of a 400 *αβ* grid cells. The high count categories of bcGST statistics supplement the singular boutique array p-values with an extra layer of interpretability on the annotation terms’ stability across a range of possible *α* and *β* values.

### 3.4 GST results are sensitive to two highly variable key ratios

If we are given a pair of *α* and *β*, then we can complete a FET contingency table using the genome or some genome-wide gene expression platform as the gene universe. However, each estimate of the *(α, β)* pair only assumes one plausible level of gene-set selection bias. By running a boutique array simulation on the TCGA data, we show that inference results are very sensitive to these two ratios and that both ratios are difficult to estimate. Hence, a single estimate of these ratios does not yield stable results. This *(α, β)* induced instability is avoided through the proposed gird-based approach in bcGST.

#### 3.4.1 Heterogeneity of *α*

First, by considering running the boutique array simulation on a set of the TCGA gene expression data, we can show that *α* is highly heterogeneous within and between different cancer datasets. Figure 4 shows the distribution of both *α* and *β* ratios for 2,219 common GO terms between 11 cancers. The parameter *α* is highly variable within individual datasets, indicated by the upper boxplot tails extending towards 1 and a large number of zeros. Despite having an overall median around 0.14 across all cancer datasets, the boxplots show that the distribution of *α* values also differ between cancers. Such variability makes estimating *α* for every GO term difficult, and therefore statistical conclusions will be more prone to errors.

**Fig. 4.**
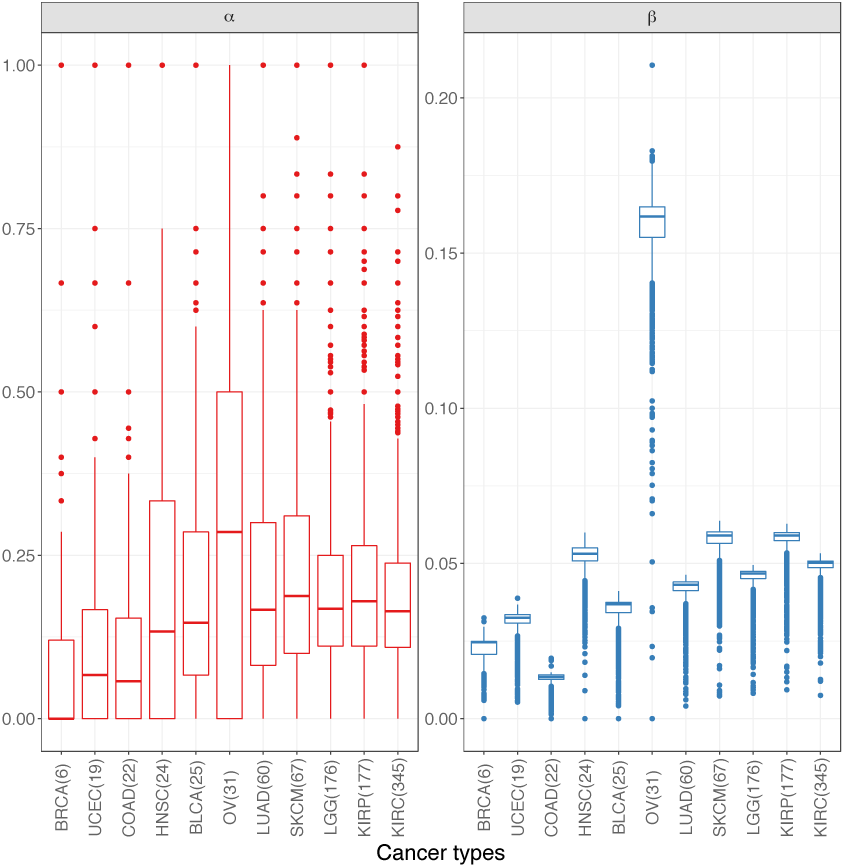
Boxplots of *α* and *β* for 11 different cancers, using a simulated boutique array on the TCGA data. The cancers are ordered by number of DE genes on the simulated boutique array. We observed high stability for *β* across different cancers (except ovarian caner) and *α* is much more heterogeneous within each cancer. This suggests the estimation of *α* will be difficult both within and between different cancers. Thus we should expect a grid-based approach to be more flexible in accommodating its instability and also robust against disease heterogeneity.

#### 3.4.2 Sensitivity of *β*

The ratio *β* also has a strong overall influence on the FET p-value. We can examine this by further studying the simulated boutique array results on the skin cutaneous melanoma (SKCM) data in TCGA.

Figure 5 is a ‘spaghetti’ plot of p-values for all GO terms. By fixing four selected *β* values, we can see the collective change of all GO terms across different values of *α*. The general trend of p-values is very sensitive to small changes in *β*, particular for smaller *α* values; which we know from Figure 4 is a particular feature of *α*. This is to be expected since even though the range of *β* is smaller than that of *α*, the construction of *β* involves the parameter *g*, which is the number of non-boutique array DE genes absent from pathways - a number which we expect to be large in applications. Small changes to this parameter will tend to produce very different estimates for other parameters.

**Fig. 5.**
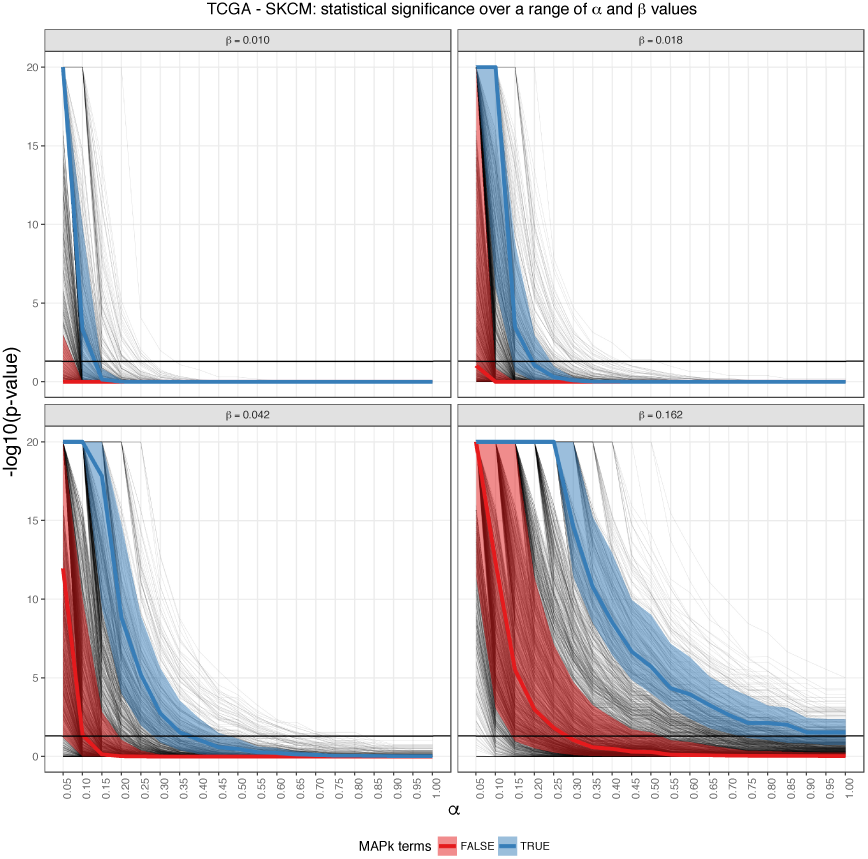
Each panel in the Figure holds a *β* value fixed. *α* is on the *x*-axis, and − log_10_ of the p-value is on the *y*-axis. Each faint black line in the diagram is a GO term, with the thick black horizontal line respond to the − log_10_ (0.05) threshold. The *β* grid points were chosen to be equally spaced over 0.01 (*β* value for COAD in TCGA simulation) and 0.16 (*β* value for OV in TCGA simulation. We note the MAP kiααnase-associated GO terms held their statistical significance across all values *α* for all values of *β*, a feature which we consider to be robust against gene set selection bias.

#### 3.4.3 Biological significance

Figure 5 contains special features that enhance biological significance as follows. First, we selected four *β* values obtained from the TCGA cancer datasets, ranging from the cancer of lowest median *β* value (COAD) to the highest median *β* value (OV). This provides us with a realistic expectation of the range of *β* in cancer studies which we could use in future visualisation heatmaps and computation of bcGST statistics. Second, over this selected range of *β*, we split the GO terms into those which are associated with MAP kinase (blue) and those which that are not (red). We computed the median of each grouped GO terms for each *α* value and also shaded the 25th percentile and the 75th percentile band for each group. Since the MAP kinase pathway is known to be associated with melanoma, we expect the GO terms associated with MAP kinase to be statistically significant across all *α*, for all selected *β* values, when compared to the rest of GO terms. We observed this trend in this plot, thus affirming these MAP kinase-associated terms retained higher their statistical significance than the rest of GO terms, and hence we can conclude these are robust against changes in both *α* and *β*. This figure allows us to see how p-values of a particular term (or a group of terms) changes with respect to other terms, under a range of possible gene-set selection bias.

### 3.5 bcGST statistic outperforms non-bias-corrected p-values as a classifier

While the bcGST grid count statistic has the ability to supplement non-bias-corrected p-values from a boutique array to recover false negatives induced by gene set selection bias on a boutique array; it can also be used as a classifier in itself to determine the significance of annotation terms. Similar to how receiver operating characteristic (ROC) curves are constructed, classification can be achieved by setting an appropriate threshold on the counts, with those pathways exceeding this threshold being judged as significant.

Figure S1 shows the ROC curve for the bcGST statistic and the FDR-adjusted FET p-values in our simulated boutique array. bcGST statistic is able to achieve similar true negative rate with a much lower false negative rate compare to simulation. This ability to reduce false negatives makes the count statistic strongly competitive against the conventional FET p-value approach on top of its robustness to bias.

### 3.6 Application to real boutique array data revealed meaningful biological results

Focusing on a comparison of AJCC tumour stage between the real melanoma NanoString data and the TCGA - SKCM data, we were able to show the value of our method. First, we can see that in practice, the concordance of p-values on two different platforms is not guaranteed to be strong as evident in Figure S2. This reaffirms the need for a correction method in the space of pathway enrichment analysis. Our grid count statistic is able to add extra insights into how stable the pathway results are. Using the same *αβ* grid of 400 on the BioCarta pathway database, we showed that, attaining high bcGST count statistic value for a number of pathways is possible despite low significance on either platform. Some of these pathways include well-known pathways such as MAPk, ERK, EGF and MET, see Table S2.

### 3.7 A Shiny-based application enable interactive visualisation of adjusted-GST results for boutique array.

The bcGST statistics is dependent on choices of:

- (*α*, *β*) grid resolution
- p-value significance threshold, and
- number of significant grid points to qualify for robustness against gene selection bias.

Providing a strict reference manual for these parameters is unrealistic and prohibitive. Hence we have built a Shiny application to assist this. Users can upload their own boutique array dataset, change and choose these parameters accordingly to visualise the effects on the final pathway analyses. The application is available at http://shiny.maths.usyd.edu.au/bcGST

## 4 Discussion

The widespread use of boutique arrays has improved sensitivity of gene expression measurements in addition to its relatively lower cost compared to platforms such as genome-wide RNA-Seq profiling. These arrays have advantages in both prediction and prognostic assessment for patients. However, researchers should also be aware of the risk of applying methodologies developed for similarly-purposed profiling platforms onto boutique arrays without justification and modification. Here, we evaluated the use of Fisher’s Exact Test, a classical tool in gene set enrichment analysis in a boutique array platform and proposed a grid count statistic that adjusts the over-representation p-value for boutique array GST interpretation.

The bcGST statistic allows us to examine the stability of p-values for each annotation term over different levels of induced gene set selection bias. For each annotation term, each count represents a statistically significant conclusion, assuming a particular level of gene-set selection bias in a boutique array experiment. A higher bcGST statistic therefore implies the statistical significance will be stable across different levels of bias, and thus the significance is less likely to be driven by the gene-set selection induced in the construction of the boutique array.

The gene set selection bias issue is not unique to boutique array platforms. The main issue in this context is that statistical tools like the FET are only interpretable with respect to the correct underlying gene universe. For example, with the mass spectrometry platform in proteomics studies, where the technology was shown to only capture up to 84% of the total annotated protein-coding genes in humans (Kim *et al*., 2014). The effects of gene set selection bias on this platform remain to be explored.

The bcGST grid count statistic depends on the number of grid points and the spacing between the grid points. Due to the heterogeneity of *α* over the interval [0, 1], choosing equal spacings is one possible natural choice. Typically, the values associated with constructing *β* is much larger in magnitude, thus we recommend to construct the grid over a more restricted interval with equal or non-linear spacing. Being aware of this issue should allow us to take note of more drastic changes in inference results. Such changes were observed in Figure 5 with comparing the *β* = 0.01 panel, where the slopes of a majority of GO terms were large relative to three other panels of *β*.

It is well demonstrated in the literature that different normalisation methods can yield wildly different and incomparable results (Law *et al*., 2014; Dillies *et al*., 2013). In practice, normalisation on a boutique array has additional analytic challenges due to the small number of genes. In the interest of generating more comparable results, we did not perform any additional normalisation on the simulated data after we subset the boutique array genes from the microarray and the RNA-seq platforms. This enables us to put a greater focus on the effect of boutique arrays as opposed to the effect of normalisation. In practice, the effect of normalisation will be combined with gene set selection bias as normalisation methods typically utilises additional information from non-expressed genes or control probes. This combination of effects will add extra challenge to adjustment methods.

## 5 Conclusion

Gene set selection bias is a significant issue with boutique array technology. Such bias is expected to affect the downstream analysis and appropriate adjustment methods are required to derive valid results. When performing gene set test on the boutique array data, we propose a grid-based evaluation technique, bcGST, along with a count statistic that is robust against the gene set selection bias, to provide interpretable visualisations which outperform a conventional p-value approach. Meaningful biological results were obtained when applied to real world data. A Shiny application is available to facilitate interactive visualisation of the results.

## Acknowledgements

We thank the Sydney Bioinformatics and Biometrics group for useful feedbacks and suggestions. The results shown here are in part based upon data generated by the TCGA Research Network (http://cancergenome.nih.gov/).

